## Supplemental Figure 1 for "Phat queens emerge fashionably late: body size and condition predict timing of spring emergence for queen bumble bees"

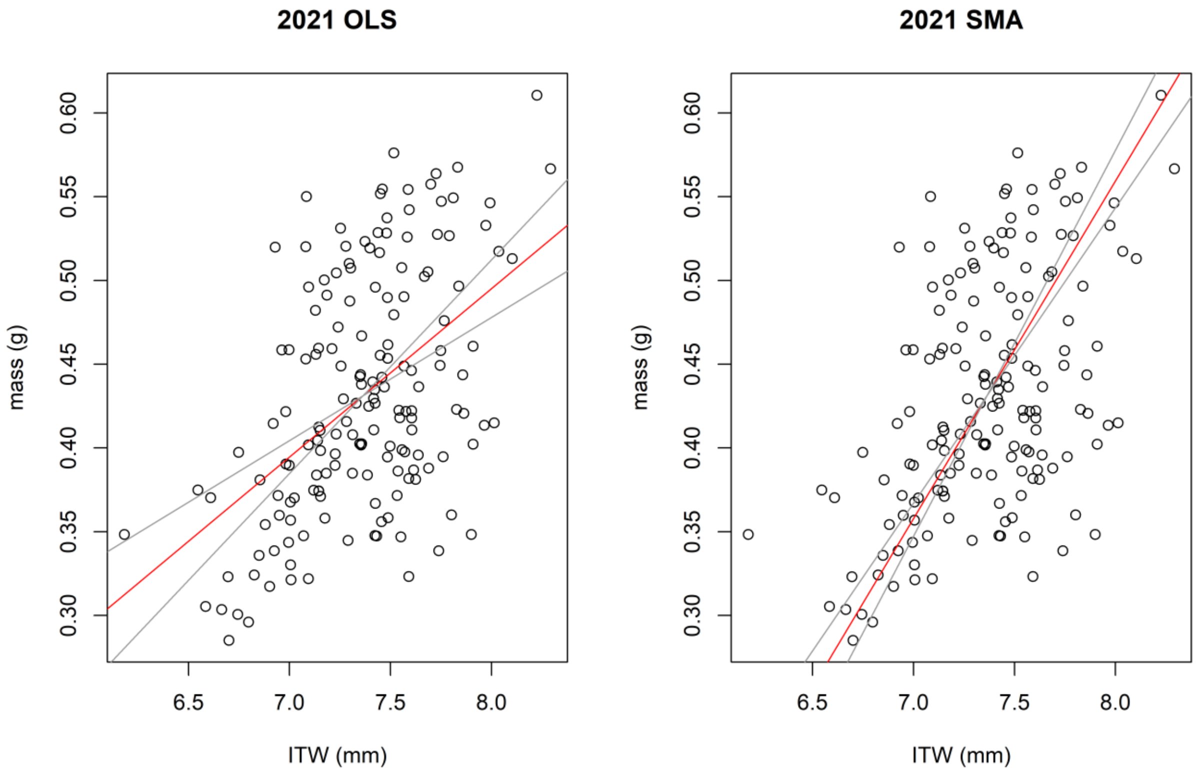


**Supplementary Figure 1**. **SMA regression was selected for 2021 after visual inspection and regression terms of mass and ITW relationships.** Body condition predictions and regression outputs were compared for 2021 using OLS (left; intercept: -0.309, slope: 0.101) and SMA (right; intercept: -1.053, slope: 0.202).
